# Short chain fatty acid butyrate promotes virus infection by repressing interferon stimulated genes

**DOI:** 10.1101/2020.02.04.934919

**Authors:** Mahesh Chemudupati, Anna C. Smith, Robert J. Fillinger, Adam D. Kenney, Lizhi Zhang, Ashley Zani, Shan-Lu Liu, Matthew Z. Anderson, Amit Sharma, Jacob S. Yount

## Abstract

Butyrate is an abundant metabolite produced by the gut microbiota and is known to modulate multiple immune system pathways and inflammatory diseases. However, studies of its effects on virus infection of cells are limited and enigmatic. We found that butyrate increases cellular infection and virus replication in influenza virus, reovirus, and human immunodeficiency virus infections. Further exploring this phenomenon, we found that addition of butyrate to cells deficient in type I interferon (IFN) signaling did not increase susceptibility to virus infection. Accordingly, we discovered that butyrate suppressed levels of specific IFN stimulated gene (ISG) products in human and mouse cells. Butyrate did not inhibit IFN-induced phosphorylation of transcription factors STAT1 and STAT2 or their translocation to the nucleus, indicating that IFN signaling was not disrupted. Rather, our data are suggestive of a role for inhibition of histone deacetylase activity by butyrate in limiting ISG induction. Global transcript analysis revealed that butyrate increases expression of more than 800 cellular genes, but represses IFN-induced expression of 60% of ISGs. Overall, we identify a new mechanism by which butyrate promotes virus infection via repression of ISGs. Our findings also add to the growing body of evidence showing that individual ISGs respond differently to type I IFN induction depending on the cellular environment, including the presence of butyrate.

**Importance:** Butyrate is a lipid produced by intestinal bacteria that can regulate inflammation throughout the body. Here we show for the first time that butyrate influences the innate antiviral immune response mediated by type I IFNs. A majority of antiviral genes induced by type I IFNs were repressed in the presence of butyrate, resulting in increased virus infection and replication in cells. This suggests that butyrate could be broadly used as a tool to increase growth of virus stocks for research and for the generation of vaccines. Our research also indicates that metabolites produced by the gut microbiome can have complex effects on cellular physiology as demonstrated by the dampening of an inflammatory innate immune pathway by butyrate resulting in a pro-viral cellular environment.

## Introduction

Of the major gut microbial metabolic end products, short chain fatty acids are of particular interest to human health. Butyrate, a 4-carbon short chain fatty acid produced from fiber metabolism, can reach concentrations as high as 140 mM in the colon, and is also present in venous blood and peripheral tissues (1, 2). Butyrate has documented roles that are largely thought to be beneficial in inflammation (3–9), adaptive immunity (2, 10–12), and in protection against bacterial infections (13, 14). Conversely, a series of classic papers showed that butyrate increased virus protein production or virion release in infections of multiple cell types with several viruses, including Epstein-Barr virus (15), measles virus (16), Borna disease virus (17), and herpes simplex virus (18). Similarly, retrovirus titers were reported to be enhanced when butyrate was added to the media of producer cells(19), leading to the use of butyrate by many laboratories in their production of retrovirus vectors. *In vivo*, butyrate and dietary fiber were shown to be protective against the influenza virus pathology in mice, despite an increase in virus titer (20). In contrast, butyrate and fiber were shown to be detrimental in the inflammatory disease caused by Chikungunya virus, a distinct RNA virus (21). The precise mechanisms by which butyrate affects viruses remain poorly understood and warrant further investigation given the ubiquity and abundance of this metabolite.

Butyrate has been known for several decades to be an inhibitor of class I and class II histone deacetylases (HDACs) (22, 23). Histone acetylation generally promotes gene transcription (24–26), and thus HDAC inhibition is suspected to be a primary mechanism by which butyrate increases retroviral vector production. Likewise, in influenza virus infection, HDAC inhibition was concluded to be a mechanism by which butyrate ameliorated disease via enhancement of adaptive immunity and by regulating tissue-damaging neutrophil numbers (20). It remains unclear why butyrate may have a opposing effects depending on the virus during infections. Further, how butyrate affects the replication of RNA viruses that do not integrate into the host genome or directly interact with histones remains unknown.

One of the most potent innate immune mechanisms against virus infections is initiated by the induction of type I interferons (IFNs), which are secreted and signal to upregulate the expression of hundreds of IFN stimulated genes (ISGs), many of which have antiviral functions (27, 28). Inborn human mutations in factors needed for production of IFNs or in the STAT1/2 transcription factors required for upregulation of ISGs are associated with high morbidity or even lethal clinical manifestations resulting from virus infections (29–33). Despite the critical role of type I IFNs in antiviral defense, the levels and activities of these IFNs are held in check by dozens of cellular proteins in order to limit their tissue damaging effects (34–39). In addition to cell-intrinsic regulatory components, recent evidence points to environmental and non-hereditary factors such as ambient temperature and humidity (40–42), as well as diet and microbiome composition (20, 43–47), in shaping antiviral immunity. Given the importance of fine-tuning the IFN response, it is likely that many regulatory mechanisms for the IFN induction and signaling pathways remain to be discovered. A more complete understanding of how these mechanisms independently influence virus infections and antiviral immune responses is needed. Here, we report a previously unknown role for butyrate in reprogramming the type I IFN response by differentially regulating the expression of more than 60% of ISGs. These results may explain longstanding mysteries regarding direct effects of butyrate on virus infections and provide new insights into our understanding of roles for butyrate in regulating disease physiology.

## Results

### Butyrate increases virus infection and replication

Given that butyrate has been reported to promote replication of several viruses, we sought to examine whether this observation held true for additional viruses relevant to human health. Given that butyrate and fiber were recently suggested to modulate inflammation during influenza virus infection (20), we first pre-treated A549 lung epithelial cells with butyrate prior to H1N1 influenza A virus infection. We observed that butyrate significantly increased susceptibility of cells to influenza virus infection as measured by percent infection via flow cytometry (Fig. 1a). We also measured infectious virus levels released in cell supernatants and found that virus titers were increased by an order of magnitude in butyrate treated cells compared to mock control cells (Fig. 1b). Since the concentration of butyrate reaches its highest level in gut tissue (1, 2), we tested whether butyrate affected susceptibility of colon cells to enteric virus infection. Indeed, we observed that reovirus infection of HT-29 colon cells and resulting virus titers were both significantly increased in the presence of butyrate (Fig. 1c and d). Likewise, since human immunodeficiency virus 1 (HIV-1) can infect and subsequently deplete gut-resident CD4^+^ T cells (48–50), we also examined whether butyrate altered HIV-1 infection of cells. Like influenza virus and reovirus, we observed a significant increase in HIV-1 infection and replicative capacity in butyrate treated THP1 monocytes compared to control monocytes (Fig. 1e and f). In sum, the net effect of butyrate on infection with three divergent RNA viruses was an increase in cellular infection and replication.

**Figure 1.**
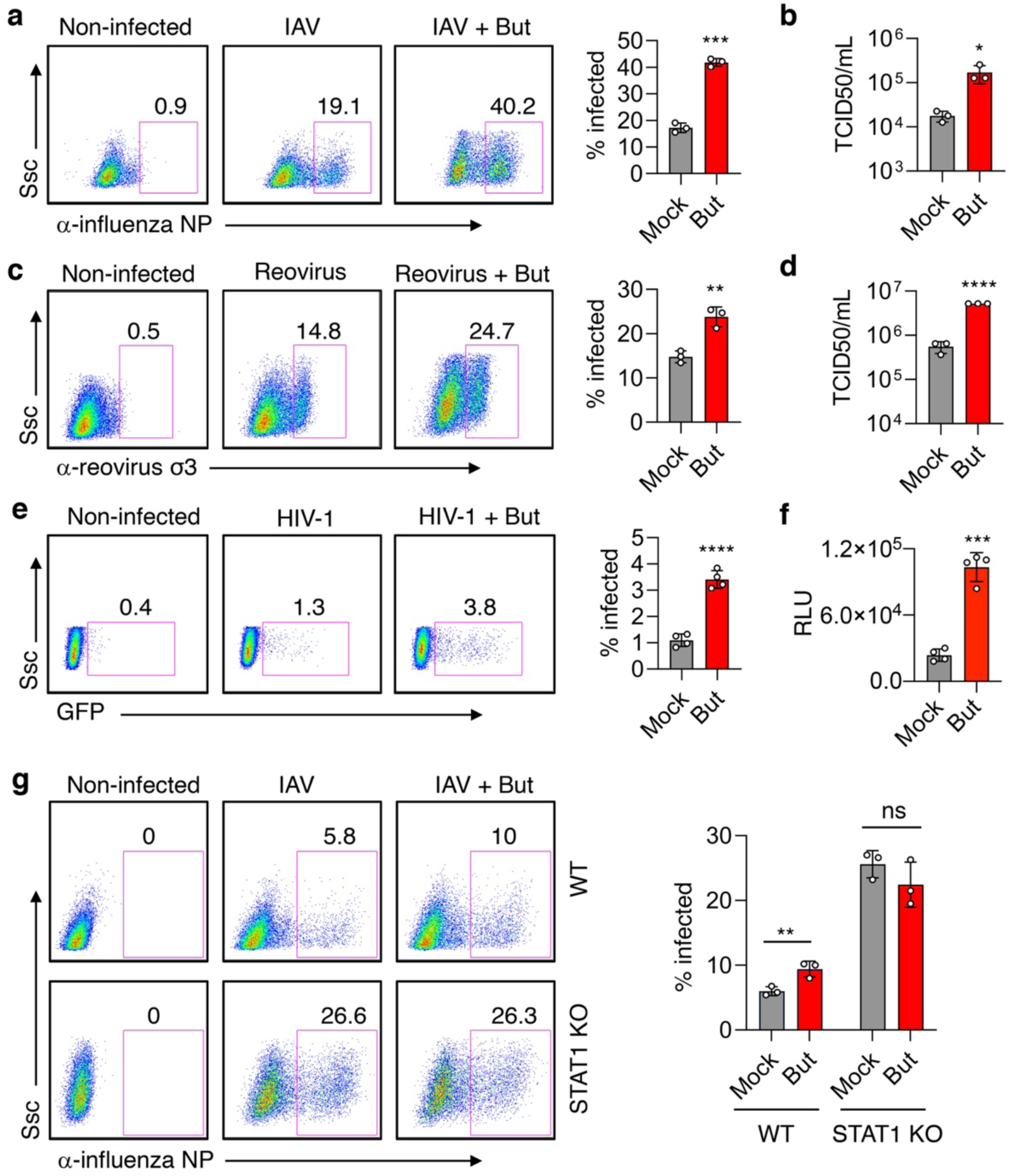
Butyrate promotes virus infection and replication. Cells were pretreated with 5 mM butyrate (But) or were mock treated for 1 h prior to infection with the indicated viruses. **A)** Left; representative flow cytometry plots from A549 cells infected overnight with influenza A virus (IAV) in the presence or absence of butyrate. Numbers indicate percentage of cells positive for IAV nucleoprotein (NP) indicating percent infection. Right; average percent infection from 4 independent infection experiments. **b)** Average virus titers in supernatants from A549 cells infected for 24 h with IAV in the presence or absence of butyrate. **c)** Left; representative flow cytometry plots from HT29 cells infected overnight with reovirus in the presence or absence of butyrate. Numbers indicate percentage of cells positive for reovirus σ3 capsid protein indicating percent infection. Right; average percent infection from 3 independent infection experiments. **d)** Average virus titers in supernatants from HT-29 cells infected for 24 h with reovirus in the presence or absence of butyrate. **e)** Left; representative flow cytometry plots from THP1 cells infected for 48 h with GFP-expressing HIV-1 in the presence or absence of butyrate. Numbers indicate the percent of cells positive for GFP indicating percent infection. Right; average percent infection from 4 independent infection experiments. **f)** Relative Luminescent Units (RLU) indicative of viral titers from from TZM-bl cells infected with cell supernatants harvested from THP1 cells that were infected with HIV-1 for 48 h in the presence or absence of butyrate. **g)** Left; representative flow cytometry plots from wildtype (WT) or STAT1 KO MEFs infected overnight with IAV in the presence or absence of butyrate. Numbers indicate percentage of cells positive for IAV NP indicating percent infection. Right; average percent infection from 3 independent infection experiments. Open circles indicate data points from independent experiments. Bars represent average values of individual data points and error bars represent standard deviation. \**P* < 0.05; \*\**P* < 0.01; \*\*\**P* < 0.001; \*\*\*\**P* < 0.0001; ns, not significant by Student’s t-test for the indicated comparisons.

We next investigated whether the increase in susceptibility to virus infection could be a result of an impaired response to type I IFNs. For this, we employed STAT1 KO mouse embryonic fibroblasts (MEFs), which are not able to upregulate IFN stimulated genes (ISGs) in response to type I IFNs. The percent infection of WT MEFs with influenza virus was reproducibly increased by 30-40% of the initial infection upon butyrate treatment (Fig. 1g), similar to results in A549 cells. In contrast, STAT1 KO MEFs, though more susceptible to infection than WT cells, were not infected at an increased rate upon butyrate treatment (Fig. 1g). Thus, intact STAT1 signaling was required for an increase in virus infection to be observed with butyrate treatment, overall suggesting that the effects of butyrate on infection may involve inhibition of STAT1 or STAT1-regulated ISGs. As an additional control, we also measured IFNβ secretion upon Sendai virus infection, a potent inducer of IFNs, and did not observe an effect of butyrate on IFNβ production (Supplemental Figure S1).

### Butyrate treatment decreases specific ISG mRNA and protein products

To test whether butyrate affects ISG induction, we measured levels of IFNβ-mediated induction of candidate ISGs in the presence of butyrate by Western blotting. We found that butyrate reduced levels of IFN-induced IFITM1, IFITM3, RIG-I, and IFIT2 in a dose-dependent manner in HT-29 colon cells (Fig. 2a). We further observed a similar effect of butyrate in reducing IFNβ-induced ISG protein levels in A549 lung cells (Fig. 2b) and RAW264.7 mouse macrophages (Fig. 2c), demonstrating conservation of the effects of butyrate on IFN responses in different cell types and in both human and mouse cells. Interestingly, we also identified that upregulation of other ISGs in response to IFN, including STAT1 and STAT2, were minimally affected by butyrate (Fig. 2b,c). The reduced levels of ISG products was not a result of butyrate toxicity as cell viability remained similar to control cells at the highest concentration of butyrate (5 mM) used in our experiments (Supplemental Figure S2). The enhanced replication of viruses in the presence of butyrate as shown in Fig. 1 further confirms that butyrate does not kill cells at this concentration.

**Figure 2.**
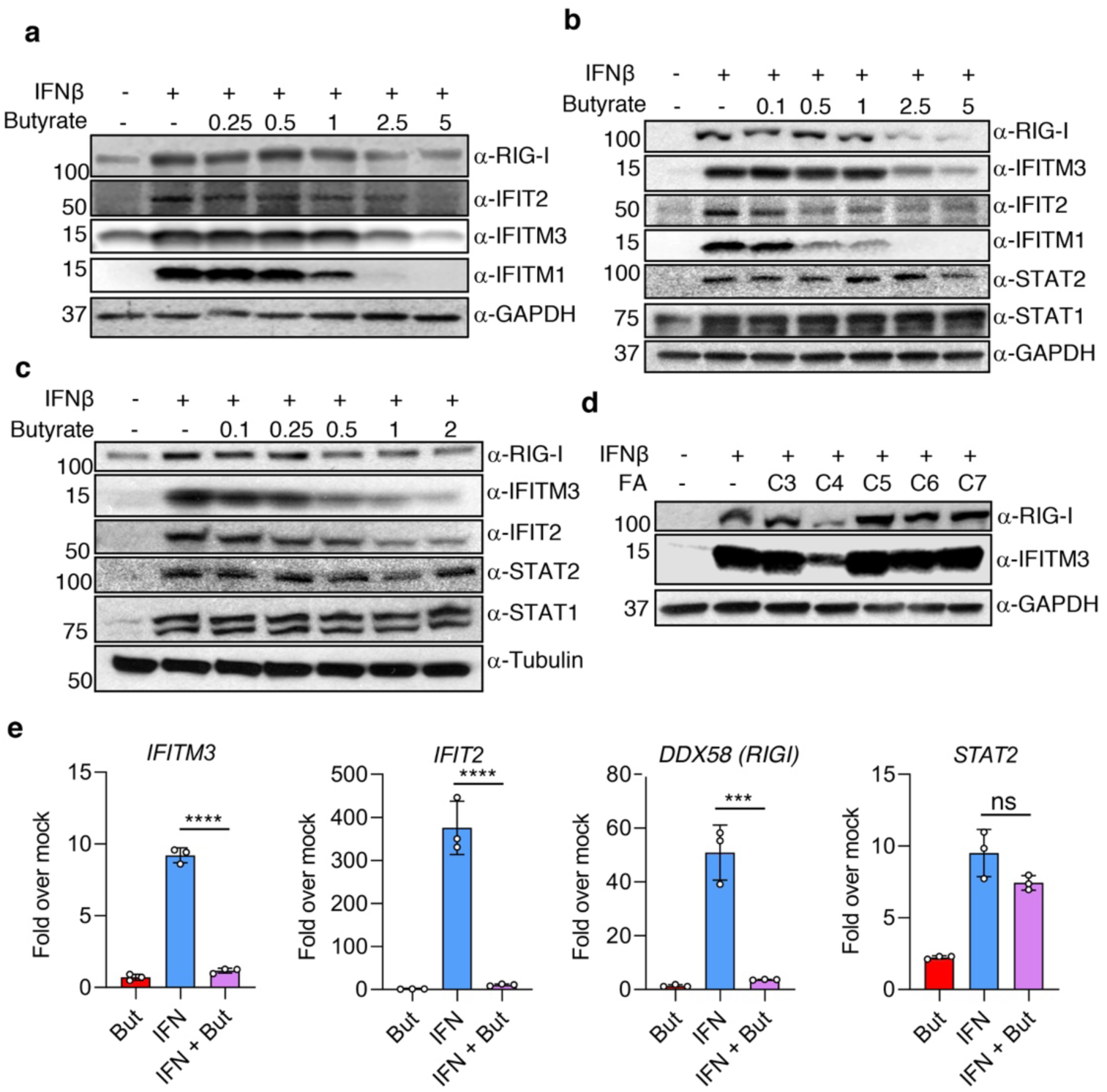
Butyrate decreases levels of a subset of ISG protein and mRNA products. **a)** HT-29 cells, **b)** A549 cells, and **c)** RAW264.7 cells were mock treated or were pretreated for 1 h with the indicated mM concentrations of butyrate followed by IFNβ or mock treatment for 24 h in the continued presence or absence of the indicated butyrate concentrations. **a-c,** Western blotting for various ISGs was performed with GAPDH and Tubulin serving as loading controls. All blots are representative of at least 4 similar experiments. **d)** A549 cells were individually mock treated or pretreated for 1 h with 2.5 mM of saturated fatty acids ranging in carbon chain length from 3 to 7 carbons (labeled C3-C7) followed by IFNβ or mock treatment for 24 h in the continued presence or absence of the indicated fatty acids. Western blotting for various ISGs was performed with GAPDH serving as a loading control. Blots are representative of two similar experiments. **e)** A549 cells were mock treated or pretreated for 1 h with 2.5 mM butyrate followed by mock or IFNβ treatment in the continued presence or absence of butyrate for 8 h. qRT-PCR was performed for the indicated ISGs. Fold change is expressed relative to mock treated cells (not treated with butyrate or IFNβ). Levels are normalized to *GAPDH* for each condition. Open circles represent triplicate measurements from a representative experiment, bars represent average values, and error bars show standard deviation. qRT-PCR data shown is representative of at least two similar experiments. Horizontal lines indicate statistical comparisons of interest. \*\*\**P* < 0.001; \*\*\*\**P* < 0.0001; ns, not significant by ANOVA followed by Tukey’s multiple comparison test.

To investigate whether other short chain fatty acids affect the cellular response to type I IFN, we pre-treated A549 cells for 1 h with a panel of saturated fatty acids of differing carbon chain lengths (three to seven carbons) prior to addition of IFNβ. We observed that, amongst our panel of fatty acids, butyrate (labeled as C4 for its carbon chain length) uniquely and potently reduced levels of the ISGs IFITM3 and RIG-I (Fig. 2d).

We next examined whether butyrate also affects the levels of ISG mRNAs by performing qRT-PCR on cells after IFNβ treatment with or without butyrate. Corroborating our Western blot results, treatment with butyrate significantly reduced the levels of *IFITM3, DDX58* (*RIGI*)*, and IFIT2* mRNA levels induced by IFN (Fig. 2e). Also, in agreement with our Western blotting, the mRNA level of *STAT2* was not significantly affected by butyrate treatment (Fig 2e). Taken together, we have discovered that butyrate suppresses the production of a subset of type I IFN effectors at the transcript level.

### Butyrate does not prevent STAT1/2 phosphorylation or nuclear translocation

Binding of type I IFNs to their cognate receptor promotes the phosphorylation of cytoplasmic STAT1 and STAT2 (38, 51). Phosphorylated STAT1/STAT2 then translocate to the nucleus where they upregulate the transcription of hundreds of ISGs. Inhibition of the phosphorylation or nuclear localization of STATs would thus prevent the transcriptional activation of ISGs. Additionally, butyrate was previously reported to inhibit type II IFN (i.e., IFNγ) signaling by preventing STAT1 phosphorylation (52, 53). We therefore assessed whether butyrate affects STAT phosphorylation or translocation to the nucleus after type I IFN stimulation. We found that STAT1 and STAT2 were phosphorylated within 15 min of IFNβ treatment in both the presence and absence of butyrate, indicating that butyrate does not inhibit STAT phosphorylation (Fig. 3a). Similarly, we observed that STAT1 and STAT2 localized to the nucleus from the cytoplasm upon IFNβ stimulation regardless of butyrate treatment (Fig. 3b). Collectively, our results demonstrate that STAT signaling downstream of the type I IFN receptor is functional in the presence of butyrate, thus identifying an inhibitory mechanism of butyrate on specific ISG induction that is distinct from the previously reported effect of butyrate on type II IFN signaling (52, 53). Further, these results are consistent with our observation that induction of some ISGs is not strongly affected by butyrate (Figure 2b, c and e).

**Figure 3.**
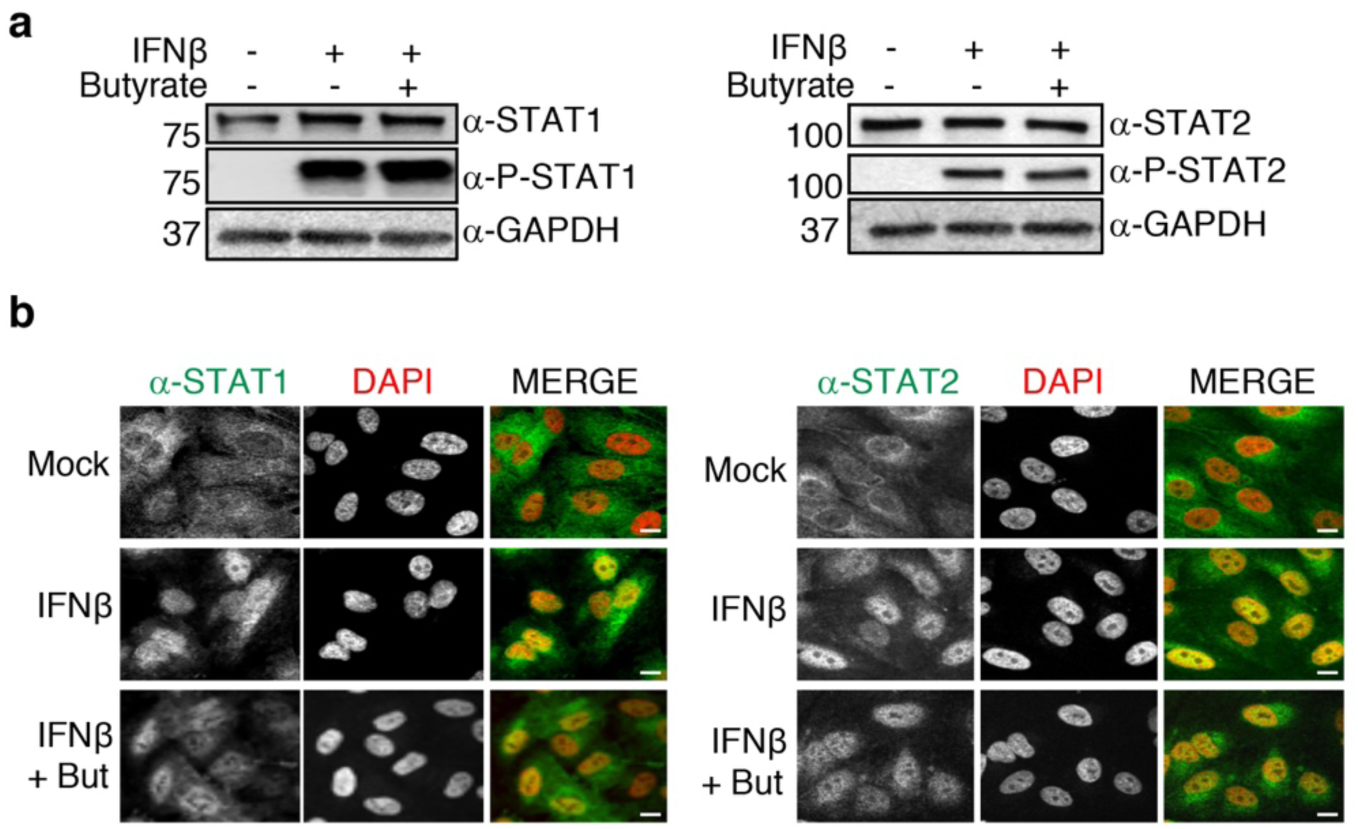
Butyrate does not prevent STAT phosphorylation or nuclear translocation upon IFNβ stimulation. **a)** A549 cells were mock treated or treated with 2.5 mM butyrate for 1 h prior to mock stimulation or IFNβ stimulation for 20 minutes. Western blotting was performed for the indicated STAT proteins and phosphorylated (P) STAT proteins with GAPDH blotting serving as a loading control. **b)** Confocal microscopy images of A549 cells treated as in **a** with staining for STAT1 and STAT2. DAPI used to visualize nuclei. Scale bar, 10 μm.

### Suppression of ISG induction by butyrate can be mimicked by other HDAC inhibitors

Since our results did not indicate an effect of butyrate on STAT signaling and translocation, its effects on ISG induction presumably result from modulation of processes downstream of STAT activity. We thus hypothesized that the known ability of butyrate to affect gene expression via HDAC inhibition may be reflected in ISG suppression by butyrate (22, 23, 54, 55). We reasoned that if butyrate is affecting ISG expression via its HDAC inhibition ability, then other HDAC inhibitors should also suppress the induction of ISGs. To test this, we pretreated A549 cells for 1 hour with a panel of HDAC inhibitors followed by IFNβ treatment. Like butyrate, treatment with the pan-HDAC inhibitor suberoylanilide hydroxamic acid (56) (SAHA) and the drug RGFP966 that inhibits the class I HDACs, HDACs 2 and 3, at the tested concentration (57, 58) both decreased levels of the ISGs RIG-I, IFITM3, and IFITM1 as measured by Western blotting (Fig. 4a). However, the class IIa HDAC-specific inhibitor TMP195 (59), and the HDAC8-specfic inhibitor 1-naphthohydroxamic acid (1-NA) (60) did not affect induction of these ISGs. Thus, chemical inhibitor studies suggest that ISG expression is at least in part under the regulation of class I HDACs, including HDAC2 and/or HDAC3, but not HDAC8 or class IIa HDACs. The decreases of ISG levels caused by butyrate, SAHA, and RGP966 treatments were accompanied by increased global histone acetylation at specific lysine residues (Supplemental Figure S3a). Similar effects of these inhibitors on ISG induction were observed in the HT-29 colon cell line (Fig. 4b).

**Figure 4.**
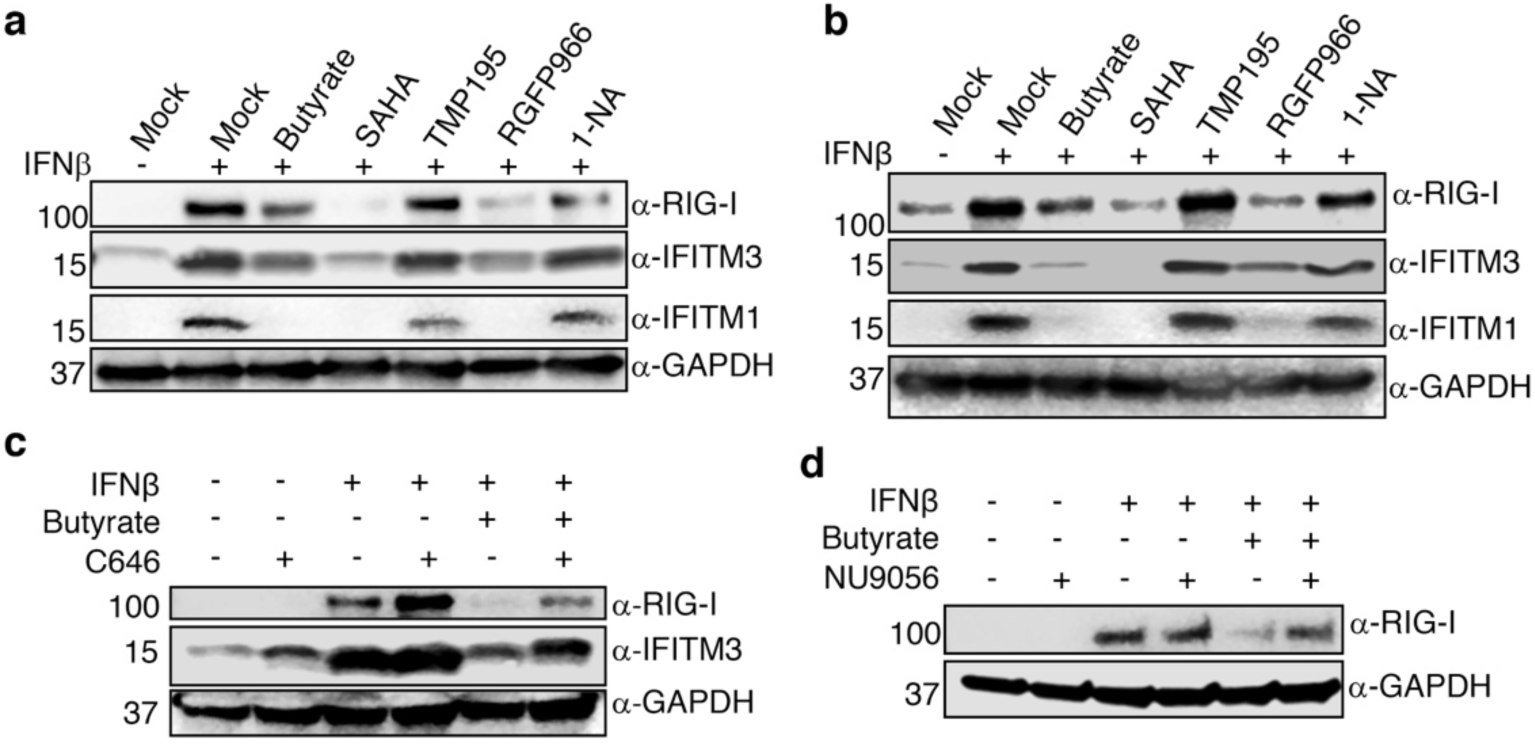
Effects of butyrate on ISGs can be mimicked by other HDAC inhibitors and can be countered by HAT inhibition. **a)** A549 cells were mock treated or pre-treated with indicated HDAC inhibitors, butyrate (2.5mM), SAHA (3 µM), TMP195 (10µM), RGFP966 (20µM), or 1-Napthohydroxamic acid (1-NA, 10µM) for 1 hour before 16 h IFNβ stimulation with continued presence of chemical inhibitors. Western blotting was performed for specific ISGs and blotting for GAPDH served as a control for loading. **b**) HT-29 cells were treated as in **a**. **c)** A549 cells were mock treated or pretreated for 2 h with HAT inhibitor C646 (10 μM), followed by addition of butyrate (2.5 mM) or vehicle control for 1 h, followed by 16 h IFNβ stimulation with continued presence of C646 and/or butyrate. Western blotting was performed for RIG-I and IFITM3 as representative ISGs and blotting for GAPDH served as a control for loading. **d)** A549 cells were mock treated or pretreated for 2 h with HAT inhibitor NU9056 (25 μM), followed by addition of butyrate (2.5 mM) or vehicle control for 1 h, followed by 16 h IFNβ stimulation with continued presence of C646 and/or butyrate. Western blotting was performed for RIG-I as a representative ISG and blotting for GAPDH served as a control for loading.

### Suppression of ISG induction by butyrate can be countered by histone acetyltransferase inhibitors

Acetylation of lysine residues on histones is mediated by histone acetyltransferases (HATs) (61, 62) while HDACs remove histone acetylation (24). Given that these two sets of enzymes mediate opposing functions, we posited that if butyrate is mediating its effect on ISGs by inhibiting HDACs, then a HAT inhibitor may counteract this effect. We thus pretreated cells with C646, a potent inhibitor of the HAT p300 (63), prior to butyrate treatment and IFN stimulation (Fig. 4c). While butyrate significantly reduced ISG protein levels upon IFN stimulation as seen previously, co-treatment with C646 partially reversed the effect of butyrate in decreasing ISG protein levels (Fig 4c). Similar results in rescuing expression of RIG-I were obtained with NU9056, a drug that inhibits KAT5/Tip60, a HAT distinct from p300 (64) (Fig. 4d). We further confirmed via Western blotting that butyrate increased the global acetylation of lysine residues on histones H3 and H4 that have been implicated in regulating gene expression, namely H3K27, H4K8, and H4K16 (61) and that this was countered by C646 co-treatment (Supplemental Figure S3b). Increased acetylation of non-histone substrates like the mRNA decay complex catalytic component CNOT7 by HATs has recently been shown to inhibit post-transcriptional gene expression by promoting degradation of mRNA (65). Thus, as an additional control, we examined knockdown of CNOT7 and found that this did not affect ISG levels, and that addition of butyrate in CNOT7*-*silenced cells remained able to decrease ISG levels (Supplemental Figure S4), suggesting that butyrate does not influence ISG levels via the mRNA decay pathway.

Our results with chemical inhibition of HDACs suggest that class I HDACs play a role in regulating ISG expression. To further explore roles of HDACs in ISG induction, we performed genetic knockdown experiments with siRNAs targeting HDACs 1, 2, and 3 either individually or in combinations to assess whether altering levels of these HDACs affects ISG induction or the effects of butyrate on ISG induction. Targeting of HDACs 1, 2, and 3 individually did not alter ISG induction by IFNβ (Supplemental Figure S5a). Targeting of the individual HDACs counteracted effects of butyrate on ISG induction to a modest, but reproducible, extent (Supplemental Figure S5a). This result was surprising in that we initially predicted that HDAC knockdown would have an additive effect with butyrate in decreasing ISG levels. However, this result may be explained by the observation that targeting of individual HDACs resulted in major compensatory feedback mechanisms in which levels of other HDACs were drastically increased (Supplemental Figure S5a). Furthermore, while certain HDAC knockdown combinations had an additive effect with butyrate-mediated inhibition of RIG-I induction, ISG levels did not correlate with levels of expression of any individual HDAC (Supplemental Figure S5b). Although compensation of HDAC levels that occurred in our knockdown experiments significantly confound interpretation of results, in conjunction with HDAC inhibitor and HAT inhibitor experiments, our results suggest that a complex interplay between multiple HDACs and HATs likely contribute to the regulation of ISG induction and the effects of butyrate.

### Butyrate has differential effects on expression of ISGs

Since butyrate can inhibit the induction of a subset of tested ISGs at the mRNA and protein level (e.g., IFITM3) but not others (e.g., STAT2), we sought to determine the extent of butyrate regulation on the global ISG transcriptomic landscape. For these experiments, total RNA sequencing (RNA-seq) was performed on mock and IFNβ-stimulated A549 cells in the presence and absence of butyrate. Global transcriptome analyses revealed that butyrate alone differentially regulated basal transcription of 882 genes, of which 821 genes were upregulated (log_2_ fold change >= 2, FDR < 0.05) and 61 were downregulated (log_2_ fold change <= −2, FDR < 0.05) (Fig. 5a and Fig. 3a, Supplemental Table 1A). The magnitude of differential expression associated with butyrate was consistent with past reports, and among the regulated genes were previously reported butyrate responsive genes such as PADI2 and CREB3L3, along with butyrate repressed genes such as AMIGO2 and DIO2 (66) (Fig 5a). Gene ontology term analysis revealed a diverse set of enriched biological associations among butyrate upregulated genes, including inflammatory and immune responses (Fig. 5b, Supplemental Table 1B), while the limited number of downregulated genes were associated with type I IFN signaling and responses to virus (Supplemental Table 1C).

**Figure 5.**
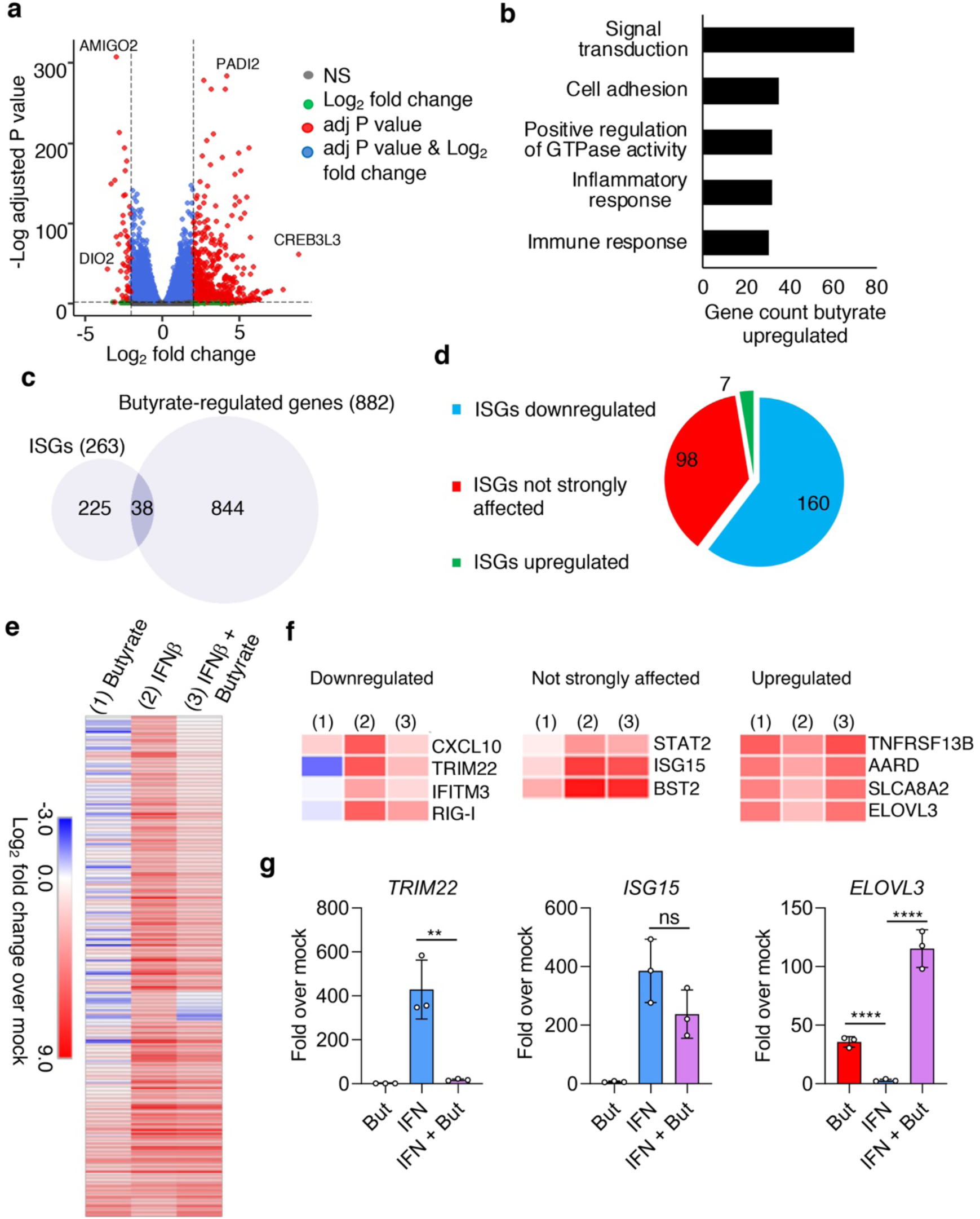
Butyrate differentially regulates baseline and IFN-induced expression of ISGs. A549 cells were mock treated or pretreated with butyrate (2.5 mM) for 1 h prior to stimulation with IFNβ or vehicle control for 8 h in the continued presence or absence of butyrate. **a-f)** RNA was extracted and subjected to RNA-seq analysis. Three biological replicates for each condition were analyzed by RNA-seq. **a)** Volcano plot of differentially expressed genes comparing mock treated and butyrate treated cells. **b)** Gene ontology term analysis of the top 5 biological processes associated with genes upregulated by butyrate. **c)** Venn diagram comparing baseline butyrate-regulated genes (4-fold or greater increase or decrease in response to butyrate treatment) to genes classified as ISGs (genes showing a 4-fold or greater increase in expression stimulated by IFNβ). **d)** Pie chart representing number of ISGs categorized based on the effect of butyrate on their mRNA levels. Downregulated genes were classified as a 4-fold or greater decrease in expression stimulated by IFNβ in the presence of butyrate. Upregulated genes were classified as a 4-fold or greater increase in expression stimulated by IFNβ in the presence of butyrate. Not strongly affected indicates all other ISGs. **e)** Heat map of all ISGs under the indicated conditions compared to mock treated cells (without butyrate or IFNβ treatment). Data represented are the mean of 3 biological replicates per condition. **f)** Magnification of specific regions in the heat map shown in **e** to visualize representative example ISGs belonging to each of the 3 ISG categories based on the effect of butyrate on their induction. **g)** Independent experimental samples were prepared as in **a-f** for validation of RNA-seq results via qRT-PCR. Fold change is expressed relative to mock treated cells (without butyrate or IFNβ treatment). Gene expression was normalized to *GAPDH* for each sample. Open circles represent triplicate samples in a single experiment. Bars represent average values and error bars represent standard deviation. Horizontal lines indicate statistical comparisons of interest. \*\*\**P* < 0.001; \*\*\*\**P* < 0.0001; ns, not significant by ANOVA followed by Tukey’s multiple comparison test.

We next examined ISG upregulation in these experiments. Our RNA-seq analysis of IFNβ-treated cells identified a total of 263 genes that were upregulated 4-fold or higher (FDR < 0.05) compared to mock treated cells, which we classified as ISGs (Supplemental Table 1D). Baseline (without IFN stimulation) expression of 38 of these ISGs was found to be upregulated or downregulated by 4-fold or greater by butyrate alone (Fig. 5c, Supplemental Table 1E). Four ISGs that included the important antiviral restriction factors IFIT2 and OAS2 were basally downregulated, while 34 ISGs including restriction factors GBP5, OASL, and BST2 were upregulated by butyrate. Thus, butyrate differentially regulates the baseline expression of several important ISGs.

We next compared the expression of ISGs in the presence of IFNβ to expression in the presence of both IFNβ and butyrate. Of the 263 ISGs, the IFNβ-induced upregulation of 160 (60%) was inhibited by 4-fold or greater by butyrate, whereas the upregulation of 96 ISGs (37%) was not strongly affected (less than a 4-fold effect) by butyrate (Fig. 5d,e; Supplemental Table 1D). Butyrate also increased the IFN-mediated induction of 7 ISGs (3%), all of which were also upregulated by butyrate treatment alone (Fig 5d,e; Supplemental Table 1D). Of the 160 ISGs that showed diminished induction in the presence of butyrate, we confirmed regulation of genes such as *IFITM3* and *DDX58* (*RIG-I*) and identified other known antiviral ISGs, such as *CXCL10* and *TRIM22* (Fig. 5f). In similar accord with our previous results, we found that butyrate did not change *STAT2* induction, and additionally discovered that induction of additional ISGs, such as *ISG15*, *BST2*, and *GBP5*, are minimally affected by butyrate (Fig 5f). The absence of a butyrate-mediated effect on IFN dependent upregulation of *BST2* and *GBP5* may be because these genes are basally upregulated by butyrate alone, as described above (Supplemental Table 1B). We also note that, consistent with our observation that butyrate did not inhibit type I IFN production (Supplemental Fig. S1), butyrate did not drastically affect induction of the transcription factor IRF7 (Supplemental Table 1D), which is an ISG that provides a feed-forward effect on type I IFN induction (67). We additionally validated RNA-seq results by performing qRT-PCR for *TRIM22*, *ISG15*, and *ELOVL3*, i.e., representative ISGs whose induction were newly found to be inhibited, unaltered, or upregulated by butyrate, respectively (Fig. 5g). Overall, we found that butyrate reprograms the magnitude of induction for at least 63% of ISGs, and have newly discovered that distinct subsets of ISGs are differentially regulated by this abundant metabolite.

## Discussion

Recent advances in our understanding of the microbiome-immunity axis has spurred interest in how gut microbiota and their metabolic products, especially short chain fatty acids, can contribute to health outcomes and potentially be used for clinical purposes (3, 68–74). In order to fully appreciate the mechanisms by which gut metabolites like butyrate exert their immunomodulatory effects, it is necessary to understand how butyrate directly influences immune programs that contribute to infection and inflammation. We report here that the short chain fatty acid butyrate can reprogram the overall type I IFN response by differentially regulating the induction of specific ISGs (Fig 5), but does not affect overall IFN production in a high MOI infection (Supplemental Figure S1). The effect of butyrate on ISG regulation does not occur at the level of STAT activation and nuclear translocation (Fig. 3), unlike the mechanism of butyrate action on the type II IFN response that was previously reported (52, 53).

Since butyrate’s inhibition of specific ISGs can be countered by HAT inhibitors (Fig. 4c,d), our findings suggest that butyrate affects ISG induction by inhibiting HDAC activity. ISG levels can also be affected by other HDAC inhibitors, including the clinically approved pan-HDAC inhibitor SAHA (56) and the HDAC2/3 inhibitor RGFP966 (57, 58) (Fig 4a,b). The effect on ISG induction by HDAC inhibitors was also accompanied by an increase in total cellular histone acetylation. Although histone acetylation is widely accepted as a posttranslational modification associated with transcriptional activation (24, 61, 62), chromatin acetylation has also been shown to be repressive in certain contexts (75, 76). Indeed, deacetylation has been shown to be required for induction of several ISGs, particularly early response ISGs, by the ISGF3 complex consisting of STAT1, STAT2, and IRF9 (35, 76–80). Therefore, it is likely that butyrate reprograms the type I IFN response by increasing histone acetylation at specific ISG loci. However, we cannot rule out the possibility that butyrate’s effects on ISGs is a result of increased acetylation of non-histone proteins, which can also be substrates of HDACs (81, 82). For example, acetylation of transcription factors has been shown to alter their affinity and subsequent transcriptional activation of target genes (82, 83).

Our findings establish that not all ISGs are induced in the same manner by the type I IFN signaling pathway. Differential induction of ISGs is suggestive of variations in trans-regulatory factors, cis-regulatory elements, local chromatin architecture, or some combination of these effects that act at individual ISG loci to control their expression. This additional level of type I IFN gene regulation could have evolved to control activation of certain ISGs that result in detrimental consequences when induced by IFN under specific environmental contexts. For example, a scenario could be envisioned in which genes that are basally upregulated by butyrate could have a deleterious synergistic effect when co-expressed with specific ISGs. A mechanism to limit the induction of such ISGs would thus confer a selective advantage under conditions of elevated butyrate concentrations.

Since it is an intestinal microbial fermentation product of dietary fiber, butyrate is present at the highest concentrations in the colon where it can be present at levels exceeding 100 mM (1, 2). The net effect of high butyrate concentration on type I IFN reprogramming in our experiments is an increase in susceptibility to infection by influenza A virus, reovirus, and HIV-1 (Fig. 1), which belong to different RNA virus families. However, it is possible that other viruses are not affected by butyrate, depending on whether butyrate suppresses induction of specific ISGs that are involved in their restriction. While high levels of butyrate could potentially elevate susceptibility to virus infections *in vivo*, butyrate has also been extensively characterized as having beneficial anti-inflammatory effects (2, 3, 6, 71, 84), which might alleviate the tissue damage resulting from viral infection and from excessive or prolonged IFN signaling. Consistent with this idea, a recent study demonstrated that mice that were fed high fiber or butyrate-rich diets had higher viral titers when challenged with influenza A virus during the early stages of infection when compared to control mice, but experienced less tissue damage to lungs during later stages of infection (20). Taken together with our findings, this suggests that high fiber diets might confer a protective advantage by reducing inflammation caused by type I IFN or other proinflammatory cytokines at the cost of temporarily increasing the overall susceptibility to virus infections.

Our findings implicate a previously unknown role for butyrate in differentially regulating type I IFN induced genes. Since butyrate can increase overall virus titers in cell cultures, butyrate treatment could be employed as an inexpensive strategy to increase the yield of viral vaccines or virus stocks that are produced in cell lines (85, 86). These results also suggest that treatment with butyrate or butyrogenic bacteria, which are being increasingly considered for therapeutic purposes (74, 87), should be evaluated in terms of a beneficial balance between anti-inflammatory and pro-viral effects.

## METHODS

### Cell Culture, interferon treatments, and drug treatments

A549, HT-29, THP1, MDCK, Vero, and RAW264.7 cells were purchased from the ATCC. TZM-bl cells were obtained from the NIH AIDS Reagent Program. WT and STAT1 KO MEFs (88) were generated by Dr. Alexander Ploss and Dr. Charles Rice (Rockefeller University). THP1 cells were gorwn in RPMI supplemented with 10% Equafetal fetal bovine serum (Atlas Biologicals). All other cells were grown DMEM supplemented with 10% Equafetal fetal bovine serum. Cells were grown at 37°C with 5% CO_2_ in a humidified incubator. Where indicated, cells were treated with human IFNβ (EMD Millipore) at a concentration of 40 units/mL. or with mouse IFNα2 (eBioscience) at a 1:1000 dilution for 24 h. Treatment with fatty acids propionate (P1386, Sigma Aldrich), butyrate (B103500, Sigma Aldrich), valerate (240370, Sigma Aldrich), hexanoic acid (21530, Sigma Aldrich), heptanoic acid (75190, Sigma Aldrich), and HDAC inhibitors Suberoylanilide hydroxamic acid (SAHA, 149647-78-9, Sigma Aldrich), TMP-195 (23242, Cayman Chemical), RGFP966 (16917, Cayman Chemical), and 1-Naphthohydroxamic Acid (6953-61-3, Sigma Aldrich) was done for 24 hours or 1 hour prior to IFN treatment, with drugs kept in culture medium for the remainder of experiments. HAT inhibitors C646 (328968-36-1, Sigma Aldrich) and NU9056 (4903, Tocris) were added to culture medium 2 hours prior to butyrate treatment, and 3 hours prior to IFN treatment, and were kept in culture medium for remainder of experiments.

### siRNA Knockdowns

Gene knockdown was achieved using Dharmacon ON-TARGET Plus Smart Pool siRNAs targeting human HDAC1 (L-003493), HDAC2 (L-003495), HDAC3 (L-003496), CNOT7 (L-012897)or nontargeting control (D-001810-10-20) with Lipofectamine RNAiMAX reagent (Life Technologies) according to the manufacturer’s protocol.

### Virus Propagation, Infection, and Flow Cytometry

Influenza virus A/Puerto Rico/8/34 (H1N1, PR8) and Sendai virus (SeV) Cantell strain were provided by Dr. Thomas Moran (Icahn School of Medicine at Mount Sinai) and stocks were propagated in 10-day-old embryonated chicken eggs (Charles River Laboratories) for 48 or 40 h, respectively, at 37°C. Influenza virus was titered on MDCK cells, and SeV was titered on Vero cells. Reovirus (Dearing strain) was purchased from the ATCC and propagated and titered using Vero cells. For influenza virus growth assays, TPCK Treated Trypsin (Worthington Biochemical) was included in A549 cell media. GFP-expressing HIV-1 pseudoviruses were generated in HEK293T cells as previously described (89). THP1 cells were treated with 2.5 mM butyrate for 1 hour prior to infection with HIV-1 pseudoviruses in duplicate wells at a multiplicity of infection (MOI) of 0.5. After 48 hours, cells were washed, fixed, and analyzed for GFP expression as performed previously^88^. THP1 cells were treated with 2.5 mM butyrate for 1 hour prior to infection with replication-competent HIV-1 at an MOI of 0.5. Cell supernatants were harvested at 48 h post-infection and virus titers were determined by infecting TZM-bl cells and measuring β-galactosidase activity using Galacto-Lite system (Applied Biosystems). For flow cytometry quantification of infection, IAV-infected cells were stained with anti-influenza NP (BEI resources, NR-19868), reovirus infected cells were stained with anti-reovirus σ3 antibody (Developmental Studies Hybridoma Bank, 4F2, deposited by Dr. Terence Dermody) and cells infected with GFP-expressing HIV were analyzed for GFP fluorescence directly. Flow cytometry was performed on a FACSCanto II flow cytometer (BD Biosciences), and analyzed using FlowJo software.

### Western Blotting and Confocal Microscopy

For Western blotting, cells were lysed with buffer containing 0.1 mM triethanolamine, 150 mM NaCl, and 1% SDS at pH 7.4 supplemented with EDTA-free Protease Inhibitor Cocktail (Roche). For phosphorylated protein Westerns, PhosSTOP (Sigma Aldridge) phosphatase inhibitor was added to lysis buffer. Primary antibodies for IFITM1 (13126, Cell Signaling Technology), IFITM3 (11714, ProteinTech), RIG-I (20566, ProteinTech), GAPDH (39-8600, Invitrogen), IFIT2 (PA3-845, ThermoScientific), STAT1 (9172, Cell Signaling Technology), STAT2 (72604, Cell Signaling Technology), pSTAT1 (7649, Cell Signaling Technology), pSTAT2 (4441, Cell Signaling Technology), Tubulin (Antibody Direct), HDAC1 (34589, Cell Signaling Technology), HDAC2 (57156, Cell Signaling Technology), HDAC3 (85057, Cell Signaling Technology), H4K16ac (13534, Cell Signaling Technology), H3K9ac (9649, Cell Signaling Technology), H3K27ac (8173, Cell Signaling Technology), H3 (4499, Cell Signaling Technology), H4K8ac (2594, Cell Signaling Technology), H4 (13919, Cell Signaling Technology) were used at 1:1000 dilutions or according to manufacturer’s protocol in both Western blotting and confocal imaging. For confocal microscopy, cells grown on cover slips were treated for 15 minutes with IFNβ, fixed for 20 min in 4% paraformaldehyde/PBS, permeabilized with 0.1% Triton X-100/PBS for 20 min, blocked with 2% FBS/PBS for 20 minutes, and consecutively labeled with primary and AlexaFluor-conjugated secondary antibodies (Life Technologies) in 0.1% Triton X-100 in PBS for 20 minutes. Cover slips were mounted on glass slides using Prolong Gold Antifade Mountant with DAPI (Life Technologies). Imaging was performed on an Olympus FluoView confocal microscope.

### Quantitative RT-PCR

RNA was extracted from A549 cells treated with DMSO (Mock) or 2.5mM butyrate, with and without IFNβ treatment for 6 hours, using the RNeasy mini kit (74104, Qiagen). cDNA was prepared from extracted RNA using the AffinityScript QPCR cDNA Synthesis Kit (600559, Agilent). PCR reactions for each sample were performed in triplicate with specific primers using iQ SYBR Green Supermix (1708887, Biorad). Relative gene expression was quantified using the 2−ΔΔCT method (90). PCR reactions were performed using the CFX96 Touch real-time system (Biorad). Normalization was performed using GAPDH levels. Primer sequences used can be found in Supplementary Table 1F.

### IFNβ ELISA

RAW264.7 macrophages were treated with 1mM butyrate for 1 hour prior to SeV infection. 24 h post infection, supernatants were collected in triplicate per condition. Mouse IFNβ concentrations from supernatants were analyzed using a DuoSet ELISA kit (DY8234-05, R&D Systems).

### RNA-seq transcriptomics and data analysis

RNA was extracted from A549 cells treated with DMSO (Mock) or 2.5 mM butyrate, with and without IFNβ treatment for 8 hours, using the RNeasy mini kit (74104, Qiagen). Three biological replicates were sequenced per experimental condition. RNA sequencing was performed at The Ohio State University Comprehensive Cancer Center Genomics Shared Resource. The mRNA libraries were generated using NEBNext Ultra II Directional RNA Library Prep Kit for Illumina (E7760L, New England Biolabs) and NEBNext Poly(A) mRNA Magnetic Isolation Module (E7490, New England Biolabs). 200 ng of total RNA (quantified using Qubit Fluorometer) was used to construct sequencing libraries. Libraries were sequenced with an Illumina HiSeq 4000 in Paired-End 150bp read mode. 17 – 20 million PF clusters (equivalent to 34 – 40 million PF paired-reads) were sequenced per sample. Each sample was inspected for quality using FastQC (http://www.bioinformatics.babraham.ac.uk/projects/fastqc). Alignment of reads was performed using Spliced Transcripts Alignment to a Reference (STAR) v.2.6.1 with human genome hg38 (91). The Bam files obtained from alignment with STAR were processed using HTSeq-count (92) to obtain the counts per gene in all samples. The read counts obtained from HTSeq-count were analyzed for differential gene expression using the DESeq2 function from DEBrowser (93) (https://debrowser.umassmed.edu/). Heatmaps were constructed using Morpheus software (https://software.broadinstitute.org/morpheus/). Volcano plots were generated with the EnhancedVolcano package from Bioconductor https://github.com/kevinblighe/EnhancedVolcano using R programming software version 3.5.3.

### Statistical analysis

Data are expressed as mean ± SD. Statistical analysis was performed using GraphPad Prism version 8.3.0 (GraphPad Software). Student’s *t*-tests were used for single comparisons between two groups. Other data were analyzed using one-way analysis of variance with Tukey’s multiple comparison test. \**P* < 0.05; \*\**P* < 0.01; \*\*\**P* < 0.001; \*\*\*\**P* < 0.0001; ns, not significant. Only statistical comparisons of direct interest to effects of butyrate are labeled and a lack of labeling does not indicate a lack of statistical significance.

## Acknowledgments

This work was supported by National Institutes of Health (NIH) grants AI130110 and AI142256 to J.S.Y, and AI25136 to A.S. M.C. and A.Z. were supported by NIH Training Grant AI112542 administered by the Ohio State University Infectious Diseases Institute. A.Z. is also supported by the National Science Foundation Graduate Research Fellowship Program. A.D.K. was supported by NIH Training Grant GM068412 administered by the Ohio State University Biomedical Sciences Graduate Program. The authors thank Ilse Hernandez for technical assistance.

---

**Supplemental Figure 1.**
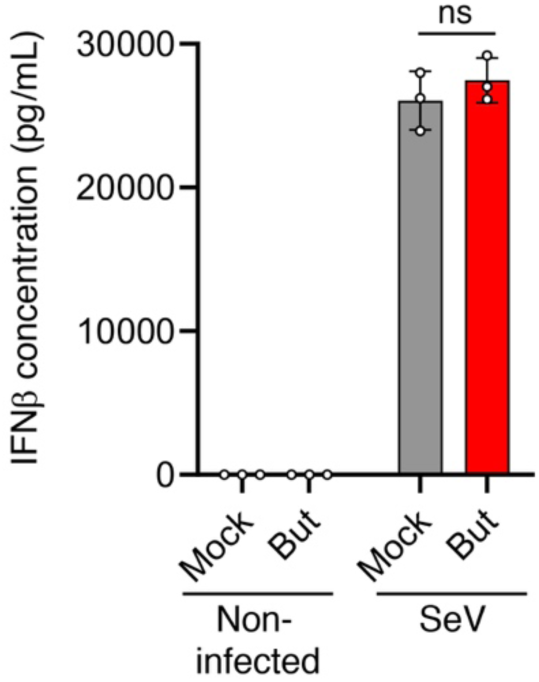
Butyrate treatment does not decrease production of type I IFN. RAW264.7 cells were mock treated or pretreated with 1 mM butyrate followed by infection with Sendai virus (SeV) at an MOI of 10 for 24 h. Supernatants were analyzed for IFNβ levels by ELISA. Open circles represent individual replicate samples from a representative experiment. Bars are average values and error bars represent standard deviation. Horizontal lines indicate statistical comparison of samples of interest. ns, not significant by ANOVA followed by Tukey’s multiple comparison test.

**Supplemental Figure 2.**
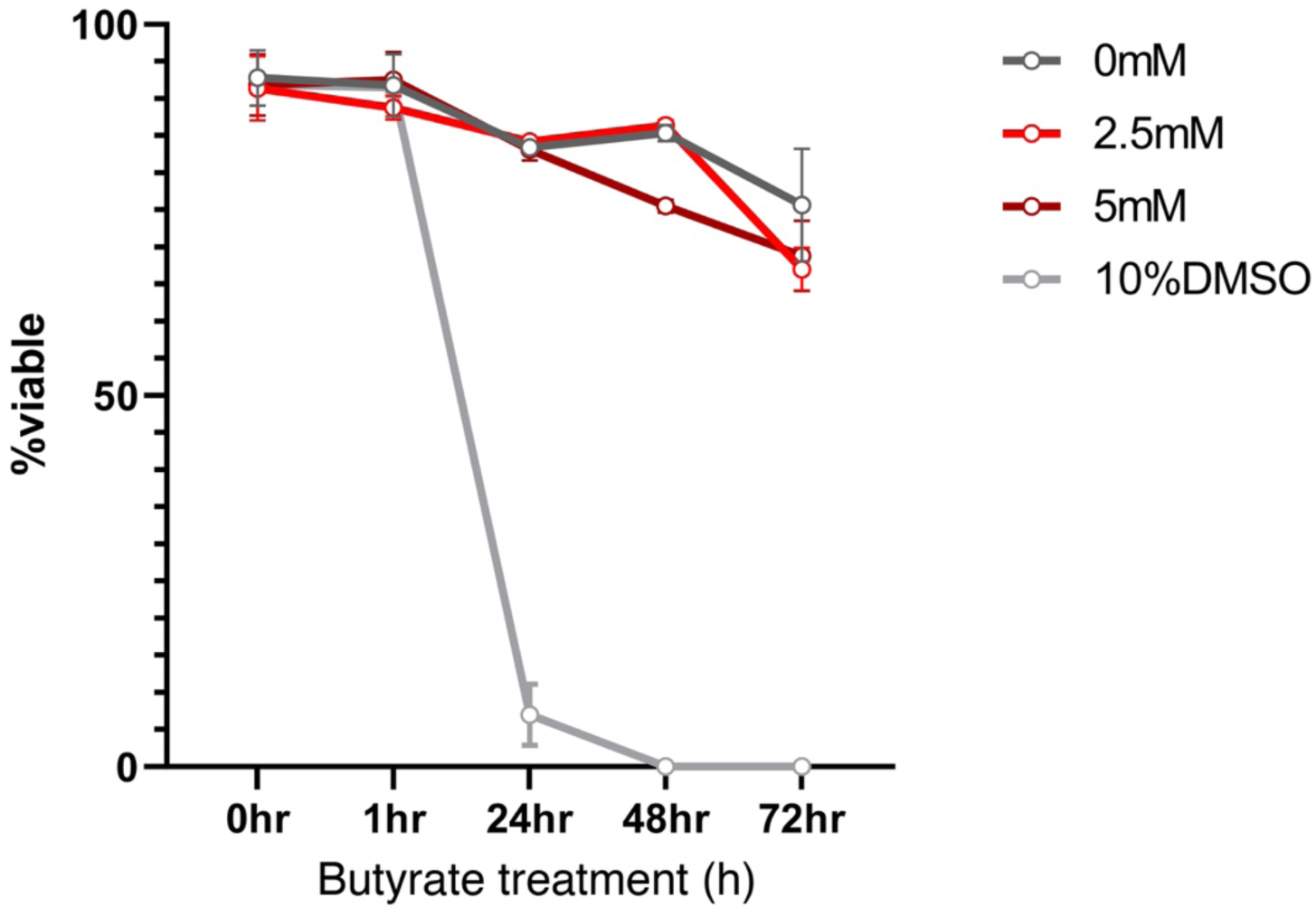
Butyrate does not reduce cell viability at 2.5 mM or 5 mM concentrations. Trypan blue staining cell viability curves for A549 cells treated with indicated doses of butyrate. 10% DMSO served as a positive control for inducing cell death.

**Supplemental Figure 3.**
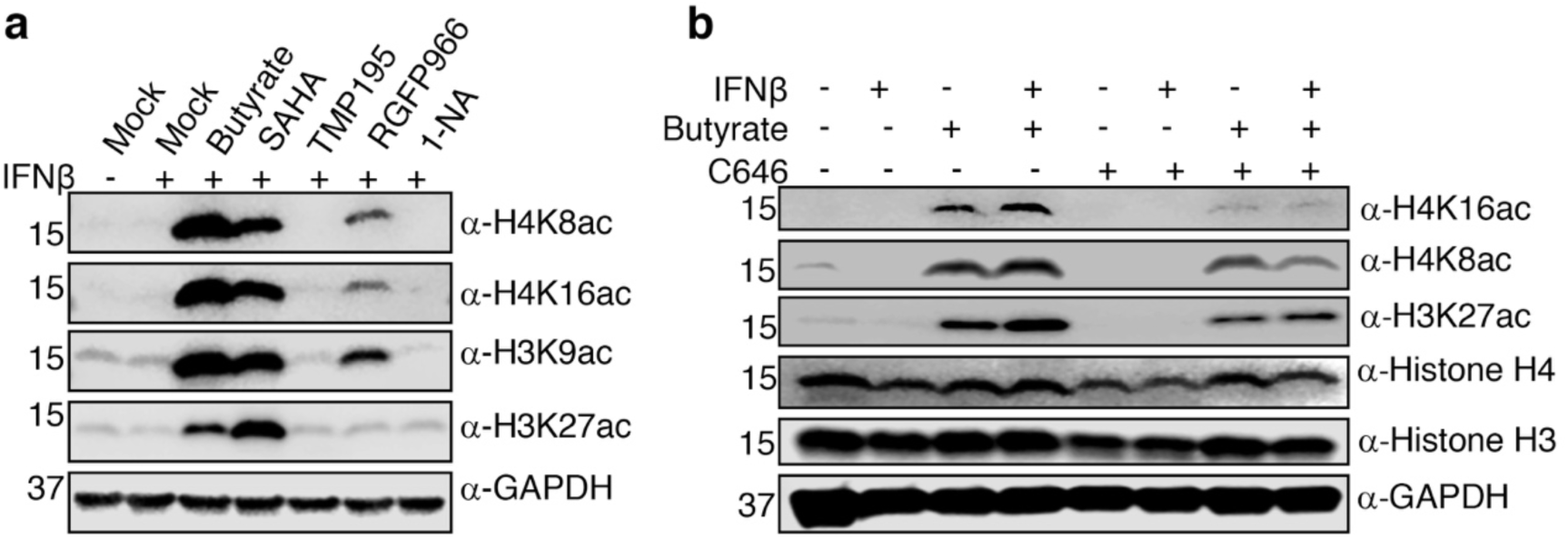
Effect of C646 and HDAC inhibitors on bulk histone acetylation. **a)** A549 cells were mock treated or pre-treated with indicated HDAC inhibitors, butyrate (2.5 mM), SAHA (3 µM), TMP195 (10µM), RGFP966 (20 µM), or 1-Napthohydroxamic acid (1-NA, 10 µM) for 1 hour before 16 h IFNβ stimulation with continued presence of chemical inhibitors. Western blotting was performed for acetylated histones and GAPDH served as a control for loading. **b)** A549 cells were mock treated or pretreated for 2 h with HAT inhibitor C646 (10 μM), followed by addition of butyrate (2.5 mM) or vehicle control for 1 h, followed by 16 h IFNβ stimulation with continued presence of C646 and/or butyrate. Western blotting was performed for histones and acetylated histones, and blotting for GAPDH served as a control for loading.

**Supplemental Figure 4.**
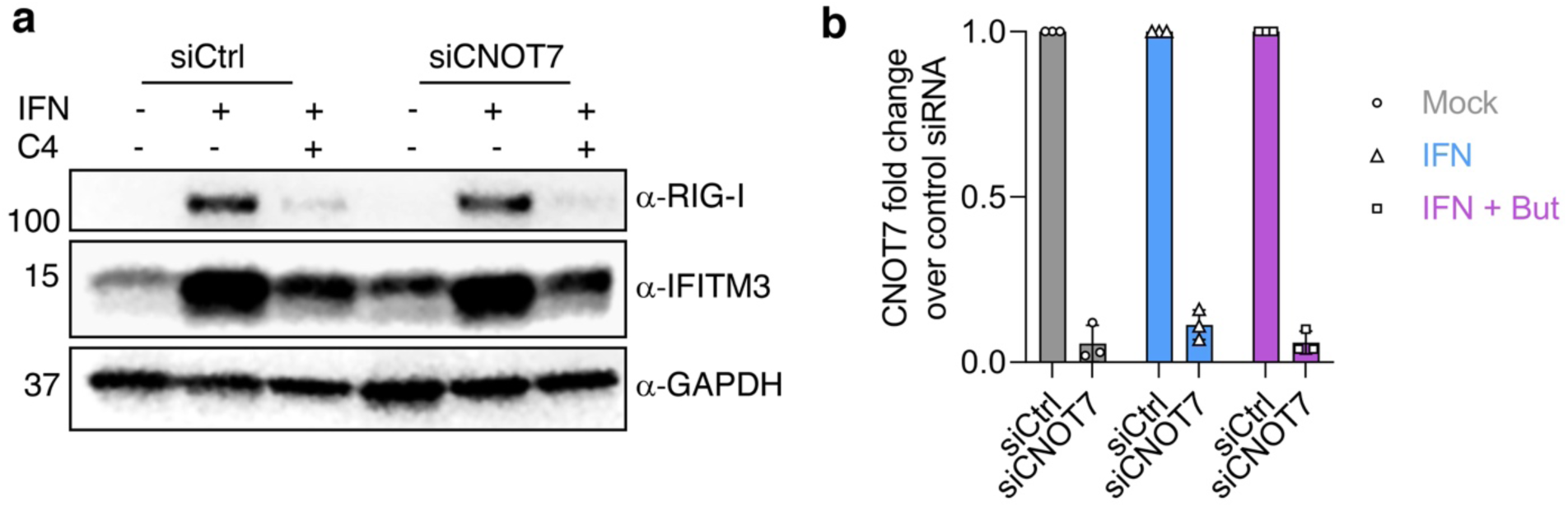
Knockdown of the CNOT7-dependent mRNA decay pathway regulator does not affect the regulation of ISG levels by butyrate. **a)** Western blot analysis of RIG-I and IFITM3 in CNOT7 knockdown (siCNOT7) compared to control siRNA (siCtrl) in the indicated conditions. **B)** qPCR analysis of CNOT7 mRNA levels in the indicated conditions verifies effective CNOT7 knockdown.

**Supplemental Figure 5.**
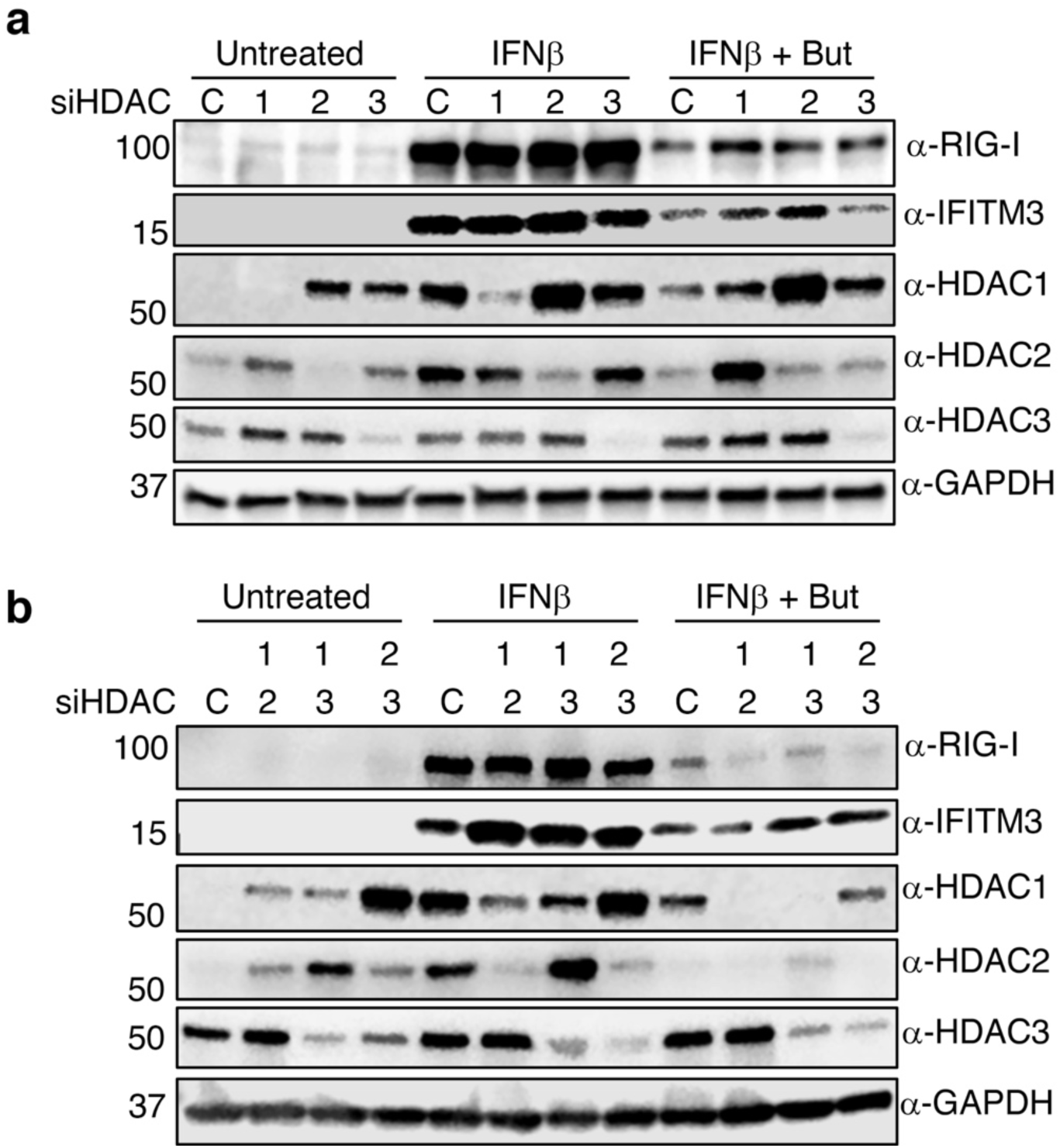
Effects of HDAC knockdown on ISG levels. siRNAs targeting HDACs 1, 2, and 3 (siHDAC) or control (C) siRNA were transfected **a)** individually or **b)** in combinations into A549 cells for 24 h prior to treatment with butyrate (But) (2.5 mM) or vehicle control for 1 h, prior to addition of IFNβ as indicated. Western blotting was performed for RIG-I and IFITM3 as representative ISGs, HDACs to examine knockdown efficiencies and expression compensation of other HDACs, and GAPDH as a control for loading.

